# Niche partitioning by resource size in the gut microbiome

**DOI:** 10.1101/2025.11.13.688124

**Authors:** Stefany Moreno-Gámez, Brent W. Anderson, Gabriel T. Vercelli, Friedrich Altmann, Andrew L. Goodman, Otto X. Cordero

## Abstract

Niche partitioning promotes diversity of the human gut microbiota. However, the molecular basis of resource specialization and niche separation in the gut remains poorly understood. Here we show that structural differences in glycan transporters drive members of the genus *Bacteroides*, common human gut commensals, to specialize on distinct chain lengths of the same fructan molecule. While species encoding canonical SusCD systems for glycan import –formed by a membrane-embedded barrel capped with a lipoprotein lid– specialized in long-chain fructans, species with smaller lidless transporters, not previously described in *Bacteroides*, specialized in short-chain fructans. Strikingly, we found that a ∼140–amino acid domain in the SusC barrel is a structural feature that governs substrate preference: deleting it does not impair transport but instead shifts uptake preferences from long-to short-chain fructans. These structural differences predict competitive outcomes *in vivo* on fructans of varying lengths, suggesting that glycan uptake mechanisms shape ecological niches in the gut and can inform fiber-based dietary interventions. Similar small lidless transporters exist across the Bacteroidota, expanding the paradigm of glycan utilization in this phylum beyond the canonical SusCD architecture.

## Main text

A key function of the gut microbiome is the degradation of complex dietary glycans that are indigestible by the human host. These glycans, collectively referred to as dietary fiber, represent a major energy source for gut bacteria and play a central role in shaping microbial community structure and metabolic activity^1–3^. Fiber molecules differ widely in their monosaccharide composition and chemical bonds, a feature that has been extensively linked to differences in the gene content of microbial glycan-degrading systems^4,5^. However, much less is known about how gut bacteria adapt to another fundamental axis of fiber chemical diversity: polymer size. Even when made up of the same monomer units, fiber molecules can differ greatly in polymerization degree, with some size distributions spanning several orders of magnitude^6–8^. Such variation is increasingly recognized as a driver of diet–microbiome interactions, influencing gut fermentation dynamics, microbial composition, and host responses to dietary interventions^9–11^. This raises the question of whether gut microbes partition niches along the fiber size spectrum, specializing on molecules of particular length, and if so, how this specialization is reflected in the evolution of their fiber utilization systems.

Here, we quantified fiber length preferences in *Bacteroides*, a dominant genus of fiber-degrading bacteria in the human gut, using inulin-type fructans. These linear chains of fructose monomers are ideal for studying length preferences, as their unbranched structure allows chain length to be varied systematically without altering other molecular features (Fig. 1a). We developed a high-throughput pipeline for growth assays in chemically defined media, allowing accurate estimation of species-specific growth rates across decreasing concentrations of each fructan (Methods). This allowed us to quantify two key physiological parameters for determining resource preferences of our species panel (Fig. 1b): the maximum growth rate at saturating resource levels (*V_max_*) and the half-saturation constant (*K_s_*), *i.e.,* the resource concentration that yields half of *V_max_*, where a smaller *K_s_* indicates higher resource affinity.

**Figure 1.**
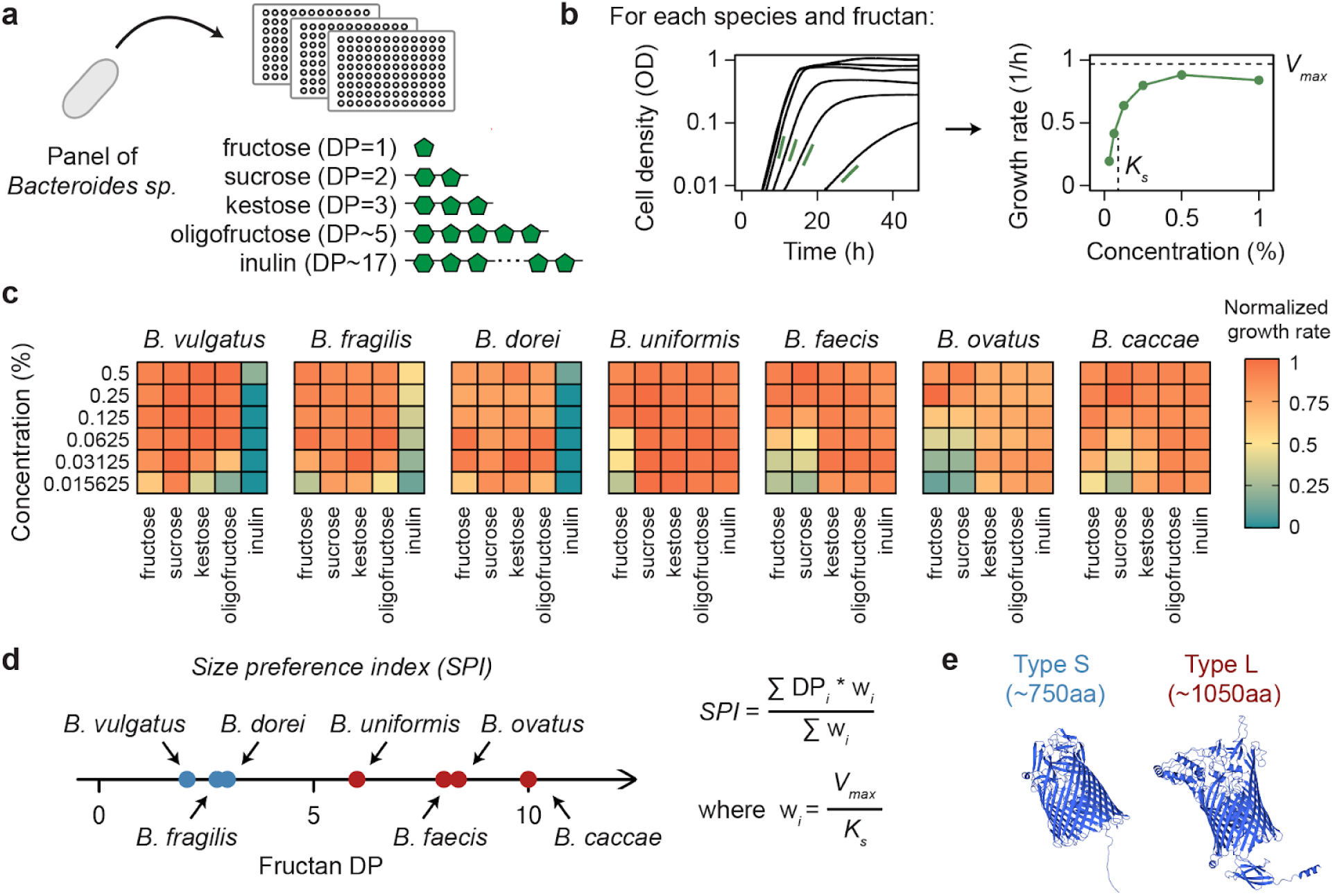
***Bacteroides* species vary in their fructan size preferences.** a) Fructan size preferences were assessed across a panel of *Bacteroides* species by measuring growth on substrates with increasing degrees of polymerization (DP). b) Monod growth parameters were estimated by culturing each species in minimal media supplemented with decreasing concentrations of each fructan type. The maximum growth rate at a saturating resource concentration (*V_max_*) and the half-saturation constant (*K_s_*) were derived from these data, with *K_s_* representing the concentration supporting growth at half of *V_max_*. Concentrations are reported as % (w/v). c) Growth rate as a function of fructan concentration and degree of polymerization (DP) for all species in the panel. For each species, growth rates were normalized to the highest observed rate across all substrates and concentrations. Species capable of maintaining high growth rates on low concentrations of short-chain fructans showed poor growth on inulin. Conversely, species proficient in inulin utilization exhibited low resource affinity for short-chain fructans. d) The size preference index (*SPI*) for every species was calculated as the average of the fructan polymerization degrees weighted by the value of *V_max_* / *K_s_* for each fructan. The *SPI* varies across species and is associated with molecular features of the fructan utilization machinery. The colors denote the type of β-barrel protein encoded by each species (panel e). e) Species specializing in long fructans encode a ∼1050 amino acid SusC-like transporter (red). In contrast, species preferring short fructans carry a smaller ∼770 amino acid β-barrel protein (blue). We refer to these as Type L and Type S, respectively.

Growth assays revealed a clear trade-off in fiber size preferences among *Bacteroides*: species capable of efficiently utilizing the longest fructan in our panel, inulin, tended to show much lower resource affinities on short fructans. In contrast, species with limited or no ability to degrade inulin performed significantly better on the shorter molecules (Fig. 1c). To quantify this trade-off more systematically, we used the values of *V_max_* and *K_s_* measured for each species in each fructan to calculate a size preference index (*SPI*) that captured the relative performance of all species across the fructan size range (Methods). This analysis showed that *Bacteroides* species differ in their *SPI* and display consistent preferences for fructans of specific polymerization degree (Fig. 1d).

Interestingly, by examining the genomic neighborhoods of the fructan-degrading enzyme family GH32, we observed a consistent association between size preferences and transporter architecture. Species specializing in longer fructans encode a SusC-like transporter of ∼1020 amino acids that acts in tandem with a SusD lid – a canonical system for fructan import that has been well characterized previously^12–14^. In contrast, species preferring short-chain fructans carry a smaller β-barrel membrane protein of ∼770 amino acids located within the same genomic region but lacking an associated lipoprotein lid (Fig. 1e). This led us to hypothesize that this smaller protein mediates short-fructan import without the aid of a SusD lid, uncovering a previously unrecognized mode of glycan acquisition in *Bacteroides*. Consistent with this hypothesis, deletion of the putative transporter gene, *BVU_1662*, in *B. vulgatus* ATCC 8482 (*Bv*) reduced growth rate and affinity of this strain for short fructans, suggesting that this β-barrel protein functions as a dedicated fructan transporter (Fig. S1). We hereafter designate the SusC-like transporter for long-fructan uptake as Type L, and the smaller lidless transporter associated with short-fructan uptake as Type S.

Genetic manipulation of *B. ovatus* ATCC 8483 (*Bo*) confirmed that the type of transporter determines fiber size preference. *Bo* was engineered by deleting the gene encoding its native Type L transporter (*BACOVA_04505*) and complementing the resulting mutant with either the same transporter or the heterologous Type S transporter from *Bv, BVU_1662* (Fig. 2a). Swapping the transporter resulted in a clear shift in fructan size preferences: while the Type L–complemented strain showed an *SPI* comparable to wild type *Bo*, the Type S–complemented strain exhibited a lower *SPI*, consistent with a preference for shorter fructans (Fig. 2b). Importantly, this shift was driven by enhanced performance of the Type S strain on short-chain fructans, including the trisaccharide kestose, relative to the Type L–complemented strain, accompanied by reduced growth on inulin (Fig. 2c and Fig. S2). Enhanced growth on short fructans was primarily due to higher resource affinity (a low value of *K_s_*) rather than to a higher maximum growth rate, which indicates that transporter type is a key determinant of size specialization, mediating a trade-off between efficient uptake of short-versus long-chain fructans (Fig. S2).

**Figure 2.**
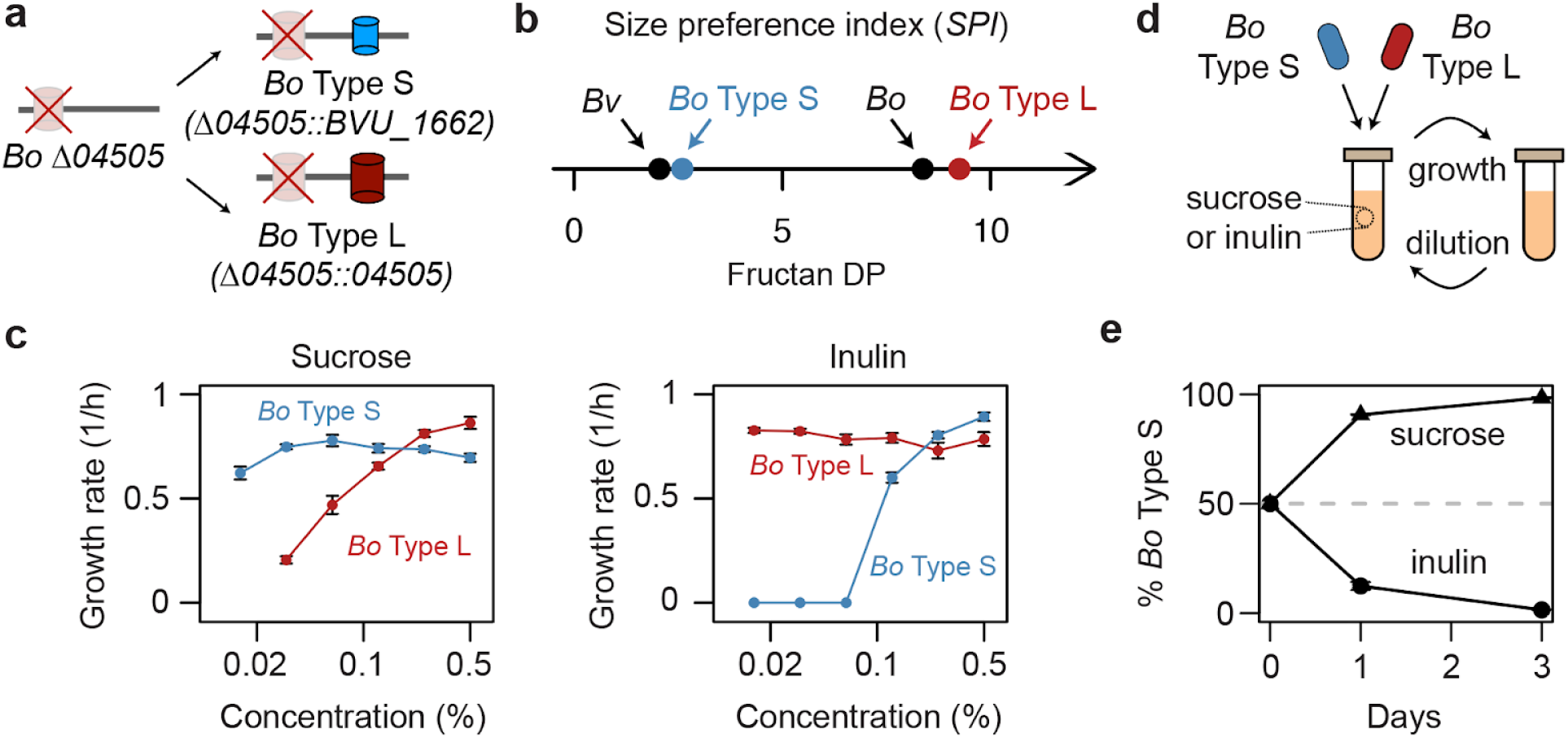
The type of transporter determines fructan size preferences and competition outcomes. a) *Bo* Type S– and Type L–complemented strains were generated by deleting the native Type L transporter (BACOVA_04505) from *Bo* and complementing the resulting mutant with either the *Bo* Type L transporter or the *Bv* Type S transporter (BVU_1662), each cloned into pNBU2 under the control of the native *BACOVA_04505* promoter. b) Swapping the native Type L transporter of *Bo* for a Type S transporter results in a change in fiber size preference from long to short fructans. c) This shift reflected enhanced performance of the *Bo* Type S–complemented strain on short-chain fructans such as sucrose, as well as fructose and kestose (Fig. S2), accompanied by reduced growth on inulin. Each data point corresponds to the mean of at least two biological replicates with error bars representing the standard error of the mean. d) Schematic of pairwise competition experiments. *Bo* strains carrying either the Type S or Type L transporter were inoculated at approximately equal initial frequencies and propagated through daily 1:100 dilutions in minimal media containing either 0.125% (w/v) sucrose or 0.125% (w/v) inulin for several transfers. e) Outcomes of competition experiments in sucrose and inulin. Each data point corresponds to the mean of three biological replicates with error bars representing the standard error of the mean. The *Bo* Type S strain outcompeted the *Bo* Type L strain in sucrose, whereas the *Bo* Type L strain outcompeted the *Bo* Type S strain in inulin, with each showing an ∼22% per-generation fitness advantage. These results demonstrate that transporter type determines competitive fitness in a substrate length–dependent manner.

Physiological differences in fiber size preferences translated into differences in competitive fitness between strains carrying different transporter types. To test this, we tagged both *Bo* complemented strains with either the Type S or Type L transporter using a system of DNA barcodes that do not confer measurable fitness effects in *Bacteroides*, and allow strain-specific quantification via qPCR^15^. We then conducted pairwise competition experiments by serially passaging the two strains in defined media containing either sucrose (short-chain fructan) or inulin (long-chain fructan) (Fig. 2d). Consistent with ecological theory, which predicts that relative fitness under resource limitation is determined by the *V_max_* / *K_s_*ratio^16^, one strain rapidly outcompeted the other depending on the substrate type. Within three transfers (∼20 generations), the Type S strain dominated in sucrose, while the Type L strain prevailed in inulin, each showing a ∼22% per-generation fitness advantage over its competitor, demonstrating that transporter architecture determines competitive fitness in a size-dependent manner (Fig. 2e).

Having established that transporter type underlies fructan size preferences, we next investigated whether structural differences between Type S and Type L transporters could account for their distinct substrate affinities. TonB-dependent transporters are defined by two core domains: a β-barrel that spans the outer membrane (Pfam family PF00593) and a plug domain (PF07715) that occludes the barrel and mediates substrate translocation into the periplasm. While we identified both domains in Type L transporters, none of these Pfam domains were annotated in Type S transporters. However, searches against other protein domain databases detected the β-barrel: CATH-Gene3D identified it as superfamily 2.40.170.20 and PANTHER identified it as model PTHR10565. Structural modelling of the Type S transporter revealed a small plug-like feature inside the barrel, but its reduced size and lack of annotation in any protein databases led us to hypothesize that the plug itself may contribute to differences in substrate preference between the two transporters.

To test this hypothesis, we deleted the plug from the native Type L transporter (BACOVA_04505), generating a mutant predicted to encode a hollow-barrel structure (Fig. 3a). Remarkably, the plug domain alone was sufficient to account for the difference in preference between long-and short-chain fructans. Growth assays revealed that the plug-less mutant exhibited a markedly lower *SPI* than wild type *Bo*, consistent with enhanced utilization of short fructans (Fig. S3). This shift resulted not only from impaired growth on inulin but also from a gain in affinity for short-chain fructans relative to wild type (Fig. 3b and S3). Because the presence of a SusD lid also distinguishes Type S and Type L transporters, we next deleted the adjacent *susD*-like gene (*BACOVA_04504*), but this did not further improve growth on short fructans (Fig. S4). Interestingly, although previous models proposed that the SusD lid maintains outer-membrane integrity when the plug is pulled into the periplasm^13^, strains lacking both plug and lid remained viable and still displayed enhanced affinity for short-chain fructans despite in principle carrying an open channel in their membrane (Fig. S4). Together, these findings establish the plug domain as a central determinant of size preference.

**Figure 3.**
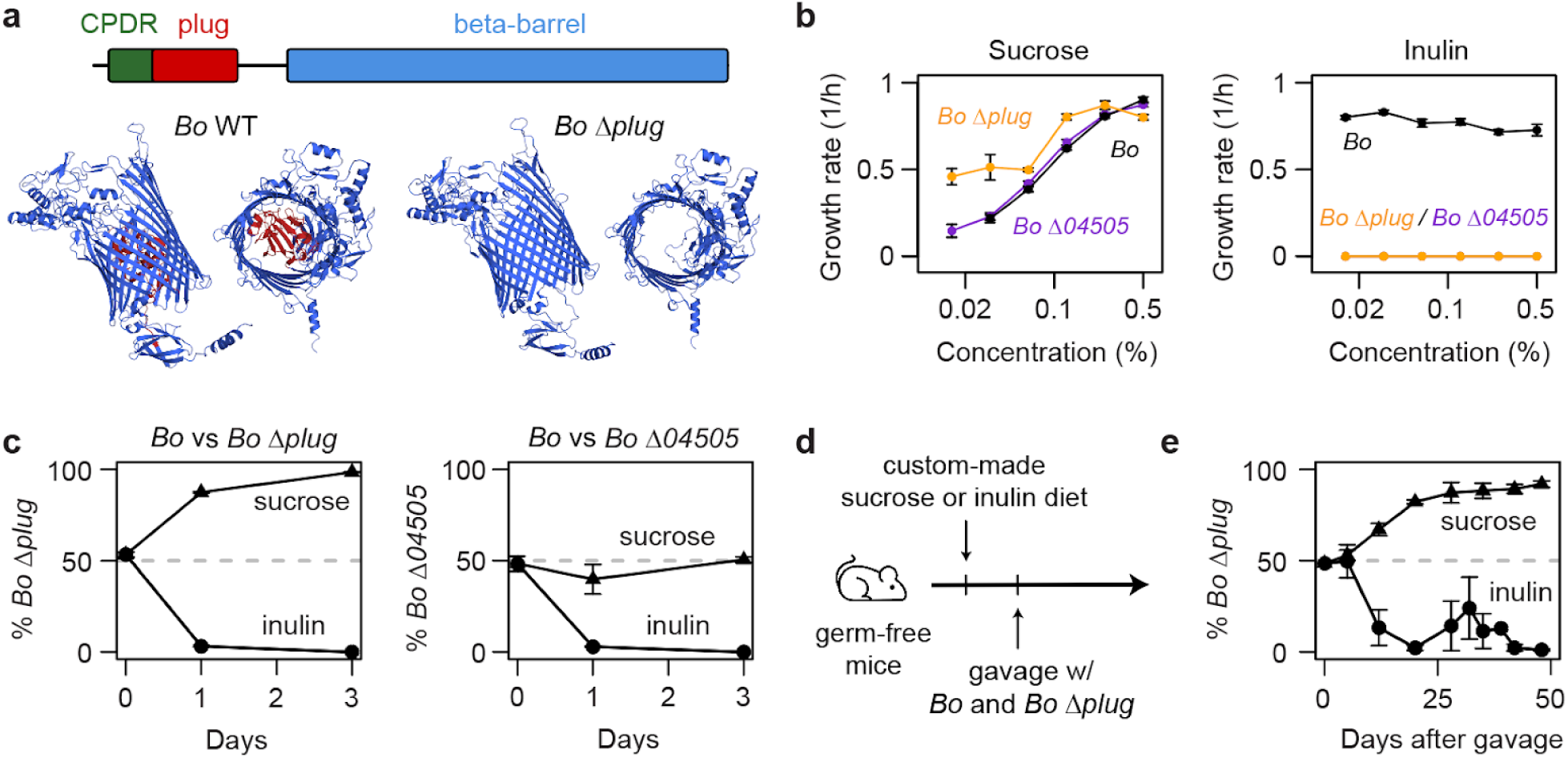
A single protein domain dictates the size preference of fructan transporters and shapes competitive fitness *in vivo*. a) (Top) Schematic of the *Bo* Type L transporter (BACOVA_04505) showing the N-terminal carboxypeptidase D regulatory-like domain (IPR008969; green), the plug domain (IPR037066; red), and the C-terminal β-barrel domain (IPR036942; blue), annotated using InterProScan. (Bottom) SWISS-MODEL structures of BACOVA_04505, generated for both the wild type and plug-deletion variants. The plug domain is highlighted in red in the wild type model. Removing the plug domain yields a hollow transporter, shortening the original protein to 885 amino acids. b) The plug-less mutant exhibits higher affinity for sucrose but loses the ability to import longer inulin chains. In contrast, a full-transporter deletion mutant shows no advantage on short fructans, indicating that the plug-less transporter remains functional. c) Affinity differences measured through growth experiments translate into distinct competitive abilities in pairwise competitions. While the plug-less mutant outcompetes wild type *Bo* in sucrose, the full-deletion mutant does not, confirming that plug removal alters size preference without inactivating the transporter. Competitions were performed by serial passaging, as described previously, with sucrose and inulin supplied at 0.125% (w/v). Each data point represents the mean of three biological replicates, and error bars indicate the standard error of the mean. d) To test these dynamics *in vivo*, germ-free mice were maintained on custom sucrose-or inulin-based diets (Table S1) for three days before being gavaged with tagged wild type and plug-less strains. Fecal samples were collected regularly over time to track strain trajectories using qPCR. e) The presence or absence of the plug domain dictates competitive outcomes *in vivo*, with sucrose or inulin diets selectively favoring plug-less or wild type strains, respectively.

Loss of the plug could in principle abolish transporter function, with enhanced growth on short-chain fructans reflecting compensatory mechanisms rather than a true change in substrate preference of the transporter. To test this, we compared the growth of the plug-less mutant to that of a strain lacking the entire transporter (*Bo* Δ*BACOVA_04505*). As expected, the full deletion mutant lost the ability to grow on inulin. However, unlike the plug-less mutant, this strain did not exhibit improved performance on short-chain fructans (Fig. 3b and S3). To further assess the ecological relevance of these findings, we conducted pairwise competition experiments using the same approach as with the complementation strains. First, we performed a competition experiment between *Bo* wild type and the plug-less mutant, which had a clear substrate-dependent outcome (Fig. 3c): while the wild type rapidly outcompeted the plug-less mutant in inulin (∼two-fold per-generation fitness advantage), the plug-less mutant consistently outgrew the wild type in sucrose (∼22% per-generation advantage). In addition, we performed a competition assay using wild type *Bo* and the full deletion mutant (*Bo* Δ*BACOVA_04505*). The results were consistent with the monoculture data: the wild type outcompeted the full deletion mutant in inulin (∼two-fold per-generation fitness advantage), but in sucrose, the deletion mutant showed no competitive advantage, and both strains remained at similar frequencies over time. These findings suggest that deletion of the plug does not inactivate the Type L transporter, but instead changes its function to favor import of short-chain fructans, leading to a shift in substrate preference.

Variation in fructan size preferences across *Bacteroides* species suggests that the relative success of individual taxa in the gut may be shaped by the chain length of fructans in the host diet. Building on our *in vitro* findings, we next asked whether the presence or absence of the plug domain would similarly affect competitive dynamics *in vivo*. To test this, we conducted pairwise competition experiments in germ-free mice colonized with the barcode-tagged *Bo* wild type and the plug-less mutant strains used for the *in vitro* competitions. Animals received sucrose-or inulin-based diets that were matched for fats, proteins, and micronutrients, following formulations from prior work (Table S1)^12,17^. Mice (*n = 4* per diet) were placed on either the sucrose or inulin diet for three days prior to bacterial gavage, and fecal samples were collected regularly to quantify strain abundances via qPCR (Fig. 3d). Consistent with our *in vitro* observations, competitive outcomes *in vivo* were diet-dependent: in mice fed the inulin diet, the wild type strain rapidly outcompeted the plug-less mutant, whereas in mice receiving the sucrose diet, the plug-less mutant became dominant (Fig. 3e). We obtained the same result in an independent experiment using separately generated mutant strains, confirming that the observed effect is robust (Fig. S5). These results indicate that the plug domain modulates competitive fitness in a substrate-dependent manner and underscore its ecological relevance in shaping niche partitioning within the gut.

Motivated by our findings that transporter architecture drives size specialization in fructans, we next investigated how broadly these transporter types are distributed across the Bacteroidota and whether they are associated with the utilization of other dietary polysaccharides beyond fructans. Existing annotations of polysaccharide utilization loci (PULs) in this phylum typically rely on the co-occurrence of SusC–SusD pairs^18–20^, an approach that would overlook Type S transporters, which lack a SusD lid and thus escape the standard PUL prediction pipeline. To overcome this limitation and capture PULs associated with both types of transporters, we developed an enzyme-centered algorithm that locates all CAZymes in a genome, retrieves their gene neighborhoods, and merges overlapping neighborhoods to generate a list of putative glycan utilization gene clusters independent of SusD annotations (Fig. S6a and Methods). From these, we selected clusters that contained a single TonB-dependent β-barrel gene predicted to be secreted by SignalP^21^.

This analysis revealed that, beyond the canonical SusCD architecture, CAZymes co-localize with TonB-dependent transporters lacking a SusD lid and, in some cases, also a plug domain, such as the Type S fructan transporters identified in this work (Fig. S6b). Remarkably, these non-canonical systems account for ∼35% of all CAZyme-associated TonB-dependent transporters identified by our pipeline. These lidless transporters likely specialize in the uptake of small substrates, since SusD is dispensable for growth on oligosaccharides in canonical SusCD systems^13,22,23^. Moreover, in these lidless transporters the plug domain is smaller or absent altogether (Fig. S6b), a feature that, as shown here, would improve affinity for small substrates. Transporters bearing a SusD lid, by contrast, almost never lack a plug domain (<1%) and are larger because they retain this domain and likely require additional extracellular loops to mediate interactions with SusD^14,24^. These structural trends result in a bimodal size distribution, indicating that the size-specialization paradigm uncovered for fructans potentially extends more broadly across TonB-dependent transporters in the Bacteroidetes (Fig. S6c). Finally, consistent with this paradigm, the number of CAZymes per cluster increases from clusters with transporters lacking both a plug and a lid to those with the classical SusCD architecture (Fig. S6b, 1.74-fold per architecture step; 95% CI= 1.69–1.80; p < 0.001). Because substrate complexity often associates with larger CAZyme repertoires^25–28^, this relationship supports the model that larger, lid-associated transporters specialize in the import of more complex or higher–molecular-weight glycans, whereas the smaller, lidless and often plugless Type S transporters are adapted for the uptake of short, simple substrates. Among the novel Type S transporters, several examples stand out as candidates for small-molecule import, including clusters predicted to target dietary compounds relevant for health such as melibiose and raffinose^29^ (Fig. S7).

Our results reveal that variation in glycan transporter architecture provides a molecular mechanism for fine-scale niche partitioning by resource size in the gut microbiome. By combining quantitative physiology, genetics, and *in vivo* competition experiments, we showed that structural features of TonB-dependent transporters, notably the presence or absence of a plug domain, tune substrate affinities (*K_s_*), thereby shifting the chain length ranges that different species can exploit. In agreement with ecological theory, relative fitness under resource limitation was predicted by the *V_max_ / K_s_* ratio, directly linking transporter structure to competition outcomes. These findings establish polymer size as a fundamental and quantifiable axis of bacterial niche differentiation in the gut, alongside monosaccharide composition, and highlight how subtle modifications in carbohydrate structure can alter ecological interactions. Beyond fructans, the bimodal distribution of β-barrel sizes and their genomic associations suggest that size-based specialization is a general principle shaping glycan utilization across the Bacteroidetes.

Together, these insights expand the paradigm of polysaccharide utilization in the Bacteroidetes by revealing that TonB-dependent transporters lacking a SusD lid, previously characterized only in other bacterial phyla^19,30–32^, also mediate glycan uptake within this group. This finding uncovers a hidden layer of specialization in carbohydrate acquisition with implications for how different members of the microbiome partition dietary resources and coexist. More broadly, our work illustrates how molecular features of nutrient acquisition systems can predict ecological interactions and competitive outcomes. Incorporating resource size into the framework of microbiome ecology will be critical for understanding how dietary fibers shape community structure and function, and may inform targeted dietary or microbial interventions to modulate the gut ecosystem.

## Methods

### Bacterial strains

Bacterial strains used in this study are listed in Table S2.

### Growth conditions and size preference index calculation

Bacteria were streaked from −80°C glycerol stocks onto Brain Heart Infusion (BHI) agar plates supplemented with 5 mg L^-1^ Hemin, 1 mg L^-1^ Vitamin K and 0.05% (w/v) L-Cysteine. For each biological replicate, a single colony was used to inoculate an overnight culture in chemically defined minimal medium containing 0.5% (w/v) glucose. The composition of this medium was based on a previously published formulation ^33^, which we modified as described in Table S3. Overnight cultures were diluted 1:1000 into minimal medium supplemented with the indicated fructan at the specified concentration. Growth was monitored under anaerobic conditions at 37°C in 96-well microplates using an Epoch plate reader (BioTek) to measure optical density at 600 nm.

Growth measurements used to estimate substrate affinities were performed at low resource concentrations and therefore tended to be noisy. To obtain reliable estimates of the maximum growth rate across all fructan concentrations, growth curves were first smoothed using a ∼3 h moving average to identify the point of fastest growth. In a less smoothed version of the same data (30 min moving average), a linear regression was fitted to the log-transformed optical density values within a window centered around that point. The width of this window corresponded to one doubling time, estimated from the growth rate obtained from the highly smoothed curve. Three biological replicates were measured for each combination of strain, fructan, and concentration. Fits were inspected manually, and curves that were too noisy at low optical densities were discarded.

To calculate *V_max_* and *K_s_* for each strain–resource combination, we used all growth rate estimates obtained across biological replicates and substrate concentrations to fit the Monod equation, *g* = *V_max_* * *S* / (*K_s_ + S*), where *S* is the fructan concentration. For some strain–resource combinations, substrate affinities could not be reliably estimated because growth rates did not decrease across the tested concentration range. In those cases, we assumed that *K_s_* was equal to the minimum *K_s_* value measured for that strain across all fructans. The size preference index (*SPI*) for each strain was calculated as the weighted average of the degree of polymerization of all fructans in our panel (fructose, sucrose, kestose, oligofructose, and inulin), where each fructan DP was weighted by its corresponding *V_max_* / *K_s_* value.

### Strain construction

Primers used in this study are listed in Table S4. DNA amplification for cloning was performed using Q5 High-Fidelity DNA Polymerase (New England Biolabs) and all genetic modifications were verified by Sanger sequencing. Gene variants and plasmids were constructed using NEBuilder® HiFi DNA Assembly. For selection and counterselection, gentamicin (200 μg/mL), erythromycin (25 μg/mL), and aTC (100 μg/mL) were added as required.

Deletions were performed using the allelic exchange vector pSIE1^34^. For each construct, 1000 bp regions flanking the target locus were cloned into pSIE1, and allelic exchange was carried out by recombination followed by counterselection using an aTC-inducible toxin, as described previously ^34^.

Complementations were performed using the pNBU2 site-specific integration vector, which integrates into the chromosome at *att* sites. In *B. vulgatus*, the transporter deletion mutant (Bv *ΔBVU_1662*) was complemented with *BVU_1662* expressed under its native upstream regulatory region (635 bp upstream of the start codon). In *B. ovatus*, the transporter deletion mutant (*Bo ΔBACOVA_04505*) was complemented with either the wild-type *B. ovatus* transporter (*BACOVA_04505*) or the wild-type *B. vulgatus* transporter (*BVU_1662*), each placed under the 288 bp upstream regulatory region from *BACOVA_04505*.

Strains were barcoded for competition experiments using pNBU2 integration vectors carrying unique 20 bp sequences^15^. Barcode sequences are listed in Table S4. For complementation strains, the pNBU2 construct used for chromosomal integration of transporter genes was modified by site-directed mutagenesis to insert the corresponding 20 bp barcode. Successful pNBU2 integration was confirmed by sequencing the att sites, and only *Bo* strains with insertion at att1 were used for all experiments.

### In vitro competition experiments

Competition experiments were performed by growing the barcoded strains overnight from single colonies in minimal media supplemented with 0.5% (w/v) glucose. Optical densities of overnight cultures were normalized to that of the strain with the lowest OD, and cultures were then diluted 1:100 into 1 mL of minimal medium containing either sucrose or inulin at 0.125% (w/v). Cultures were grown for 24 h and then diluted 1:100 into fresh medium with the same composition. This dilution–growth cycle was repeated for four consecutive days.

DNA was extracted from cultures after every transfer, and relative abundances were quantified by qPCR using a CFX96 instrument (Bio-Rad) and the Luna Universal qPCR Master Mix (New England Biolabs).

### Mouse experiments

All mouse experiments were performed using protocols approved by the Yale University Institutional Animal Care and Use Committee. Germ-free C57BL/6 mice (8-12 weeks old) were singly-housed in cages within flexible plastic gnotobiotic isolators and were maintained on a 12h light/dark cycle. Mice were maintained on an autoclaved standard diet (5K67 LabDiet, Purina) *ad libitum* and then switched to an irradiated custom diet from Bio-Serv (also *ad libitum*) three days before gavage (Table S1). Mice on the inulin diet were maintained in a separate isolator from mice on the sucrose diet, and bedding was changed after switching from standard diet to custom diet to prevent coprophagy. For colonization, tagged *Bo* wild type and *Bo* Δplug (BACOVA_04505) strains were streaked on BHI agar supplemented with 10% horse blood (Quad Five), gentamicin, and erythromycin. Single colonies were inoculated into TYG medium^35^ and grown anaerobically at 37°C for 16-20 h. Cultures were centrifuged and resuspended in sterile PBS. The OD of each strain was normalized to 1 in PBS, the strains were mixed 1:1 by volume, and 200 μL was administered by oral gavage per mouse.

Fecal pellets were collected weekly and DNA extraction was performed using phenol-chloroform as described previously^36^. Relative frequencies for both strains were measured using qPCR in the same way as for the *in vitro* competitions.

### Bioinformatic and computational analyses

Reference genomes for all species in the Bacteroidota phylum (NCBI Tax ID 976) were obtained from RefSeq using the NCBI Datasets command-line tool (CLI). We retained only reference assemblies at scaffold level or higher, yielding 1,496 genomes. Protein-coding genes were predicted using Prodigal^37^ in single-genome mode (-p single-m). CAZyme annotation was performed with dbCAN^38^ using the default thresholds and only HMMER-based hits were retained.

Putative glycan utilization clusters were identified by searching each genome for CAZyme annotations in the GH, PL, CE, and CBM classes, which encompass polysaccharide-degrading and carbohydrate-binding functions. For each identified gene, a local genomic neighborhood of ±3 genes was defined. Neighborhoods with overlapping genes were then merged, yielding a non-redundant set of putative glycan utilization clusters for each genome. For each cluster, only those in which exactly one member protein contained a TonB-dependent barrel domain were retained, as determined by hmmsearch hits to the Pfam^39^ profile PF00593 or the PANTHER^40^ profile PTHR32552. The PANTHER model was included because PF00593 did not consistently detect the shortened barrels characteristic of Type S transporters (e.g., *B. vulgatus BVU_1662*). Clusters were further filtered to retain only those in which the TonB-dependent barrel protein carried an export signal (SP, LIPO, TAT, TATLIPO, or PILIN) as predicted by SignalP ^21^. The TonB-dependent barrel proteins were then classified based on the presence of a plug domain (Pfam model PF07715), and the ±2 gene neighborhood surrounding each barrel was examined for proteins with SusD-like domains (Pfam models PF07980, PF12741, PF12771, PF14322), resulting in the three categories shown in Fig. S6. HMM searches were run with gathering thresholds when defined, otherwise with a domain E-value cutoff of 1e-3.

Docking simulations of melibiose and raffinose to the putative transporter from *B. oleiciplenus* were performed using CB-Dock2^41^, which performs blind docking guided by automatically detected surface cavities.

## Acknowledgments

We thank Eric Martens for providing *Bacteroides dorei* DSMZ 17855 and Beneo-Orafti for supplying the inulin and oligofructose used in this study. We are grateful to Hera Vlamakis and Ramnik J. Xavier for providing bacterial strains from the BIO-ML cohort. We also thank Hera Vlamakis, Akshit Goyal, and past and current members of the Cordero lab for valuable discussions. S.M-G thanks the Dutch Research Council (NWO) for support through the Rubicon fellowship (019.201EN.041), the James S. McDonnell Foundation for support through the Understanding Dynamic & Multi-Scale Systems postdoctoral fellowship and the Wissenschaftskolleg zu Berlin. This work was supported by NIH grants R35GM118159 and R01DK133798 to A.L.G.

## Supplementary Tables

Table S1. Composition of rodent diets manufactured by Bio-Serv. All units are in grams.

Table S2. List of bacterial strains used in this study.

Table S3. Composition of chemically defined minimal medium. Table S4. List of primers used in this study.

## Supplementary Figures

**Figure S1.**
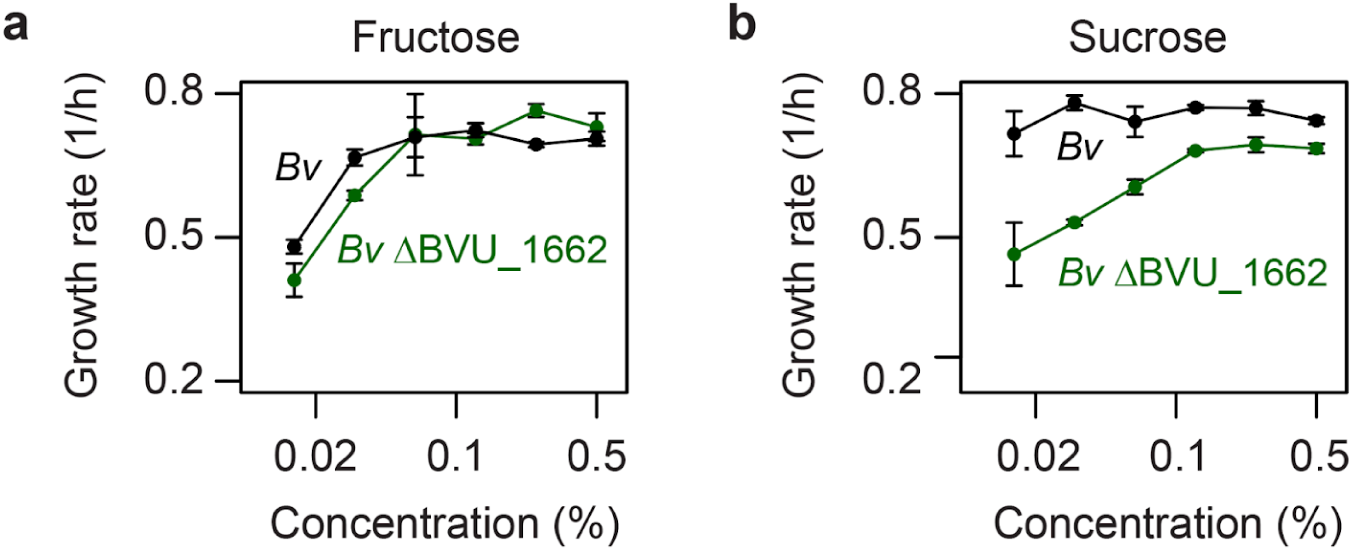
Deletion of the putative transporter gene, *BVU_1662*, from *Bv* leads to lower substrate affinities in a) fructose and b) sucrose, indicating that this gene is used for uptake of short fructans. Each data point corresponds to the mean of at least two biological replicates with error bars representing the standard error of the mean.

**Figure S2.**
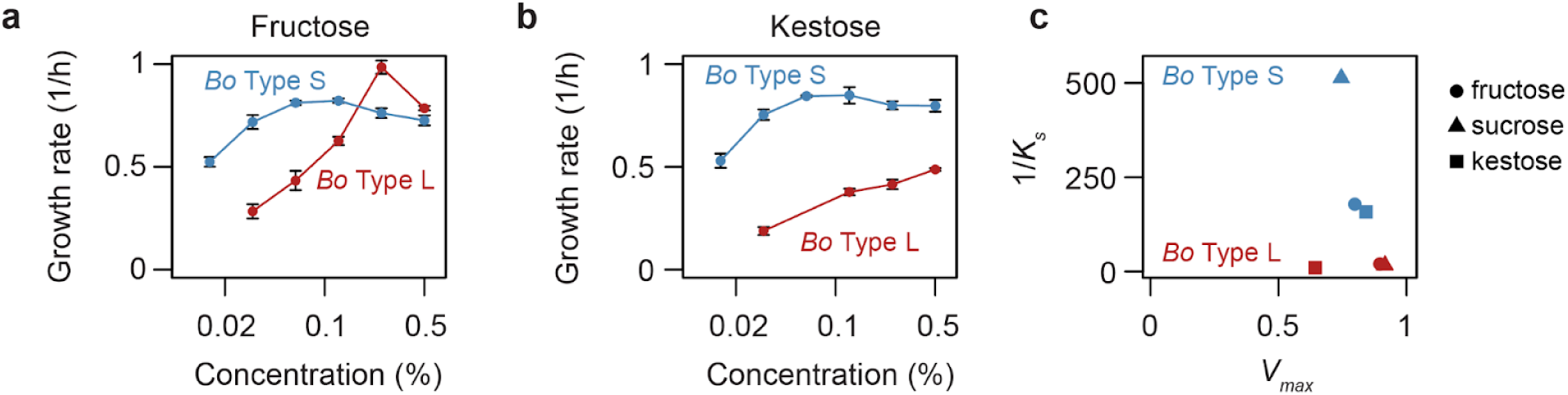
The *Bo* Type S-complemented strain (*Bo ΔBACOVA_04505*::*BVU_1662)* has a better growth performance than the *Bo* Type L-complemented strain (*Bo ΔBACOVA_04505*::*BACOVA_04505)* in a) fructose and b) kestose, indicating that the Type S transporter is more efficient at importing short fructans than the Type L transporter. Each data point corresponds to the mean of at least two biological replicates with error bars representing the standard error of the mean. c) Improved growth performance on short fructans was primarily driven by higher substrate affinity rather than by an increased maximum growth rate. Strain identity is indicated by color and resource type by shape.

**Figure S3.**
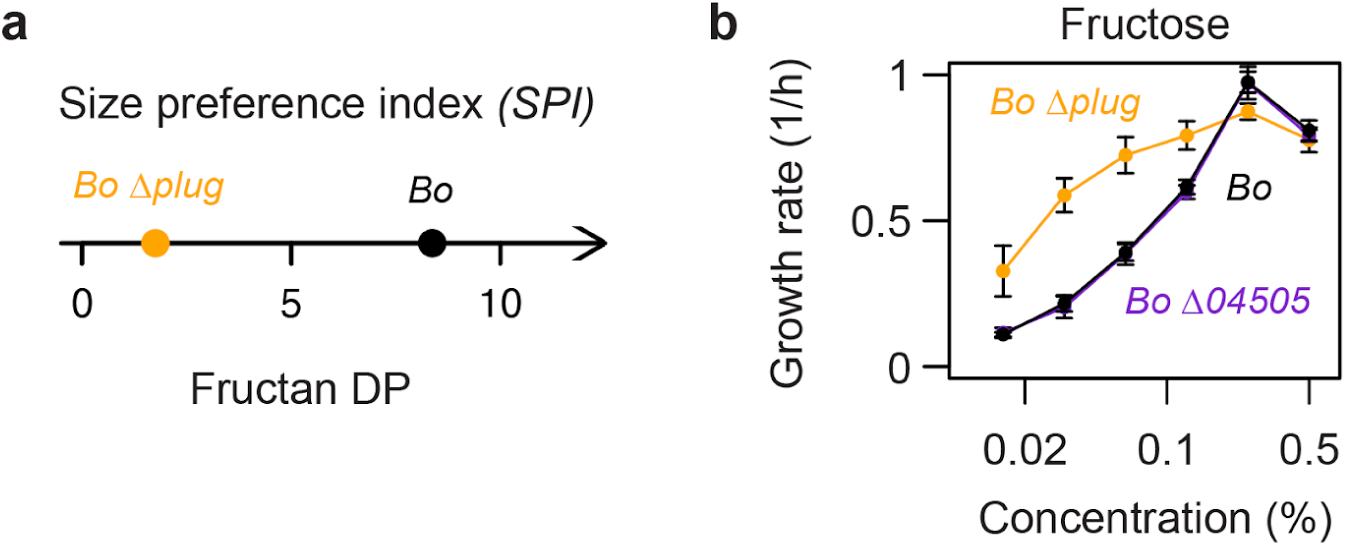
a) Deleting the plug domain from the Type L transporter, BACOVA_04505, leads to a shift in fructan size preference towards shorter molecules. b) The plug-less *Bo* mutant has higher substrate affinity for fructose than wild type *Bo* and the full transporter deletion mutant, indicating that the transporter remains functional after plug domain deletion. Each data point corresponds to the mean of at least two biological replicates with error bars representing the standard error of the mean.

**Figure S4.**
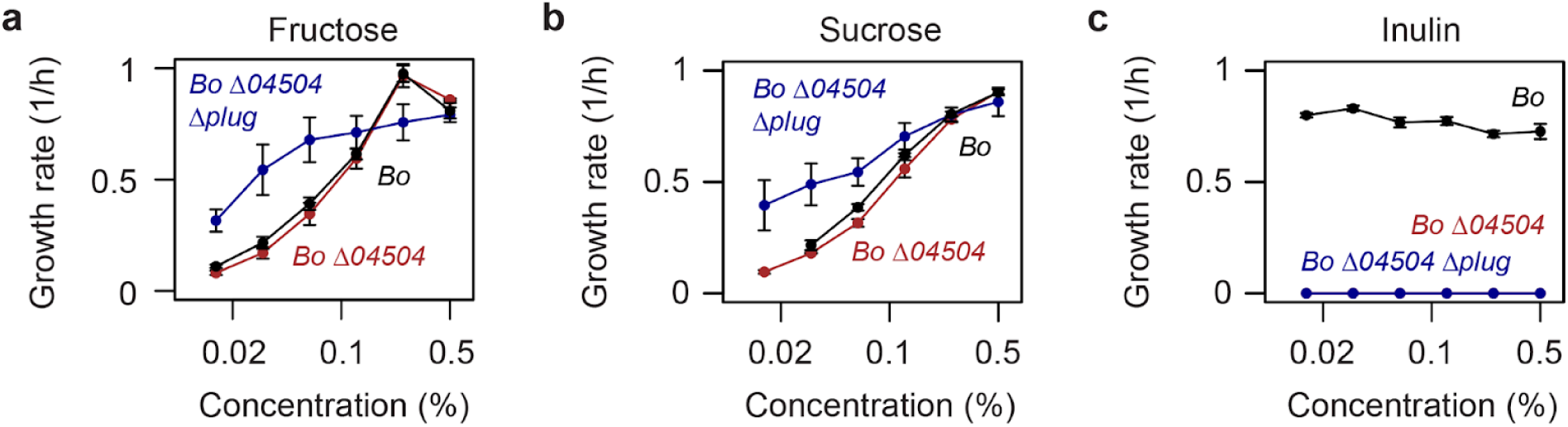
Deleting the SusD lipoprotein lid, BACOVA_04504, did not lead to improved affinity for either a) fructose or b) sucrose, but only resulted in impaired growth in c) inulin. This shows that the absence of a SusD lid does not enhance short-fructan import by Type S transporters, and reaffirms that a SusD lid is required for inulin uptake. Notably, when the plug domain was deleted in addition to SusD, the resulting strain displayed higher affinity for short fructans than both wild type *Bo* and *Bo ΔBACOVA_04504*. This confirms the plug as a critical determinant of size preference and further shows that strains lacking both the plug domain and the SusD lid remain viable. Each data point corresponds to the mean of at least two biological replicates with error bars representing the standard error of the mean.

**Figure S5.**
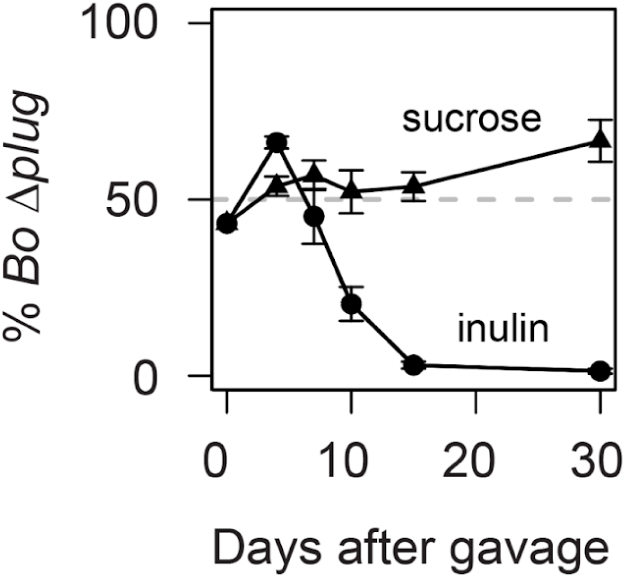
Replicate mouse competition experiment performed with wild type *Bo* and an independently generated *Bo Δplug* strain distinct from that used in Figure 3. Each data point corresponds to the mean of four biological replicates with error bars representing the standard error of the mean.

**Figure S6.**
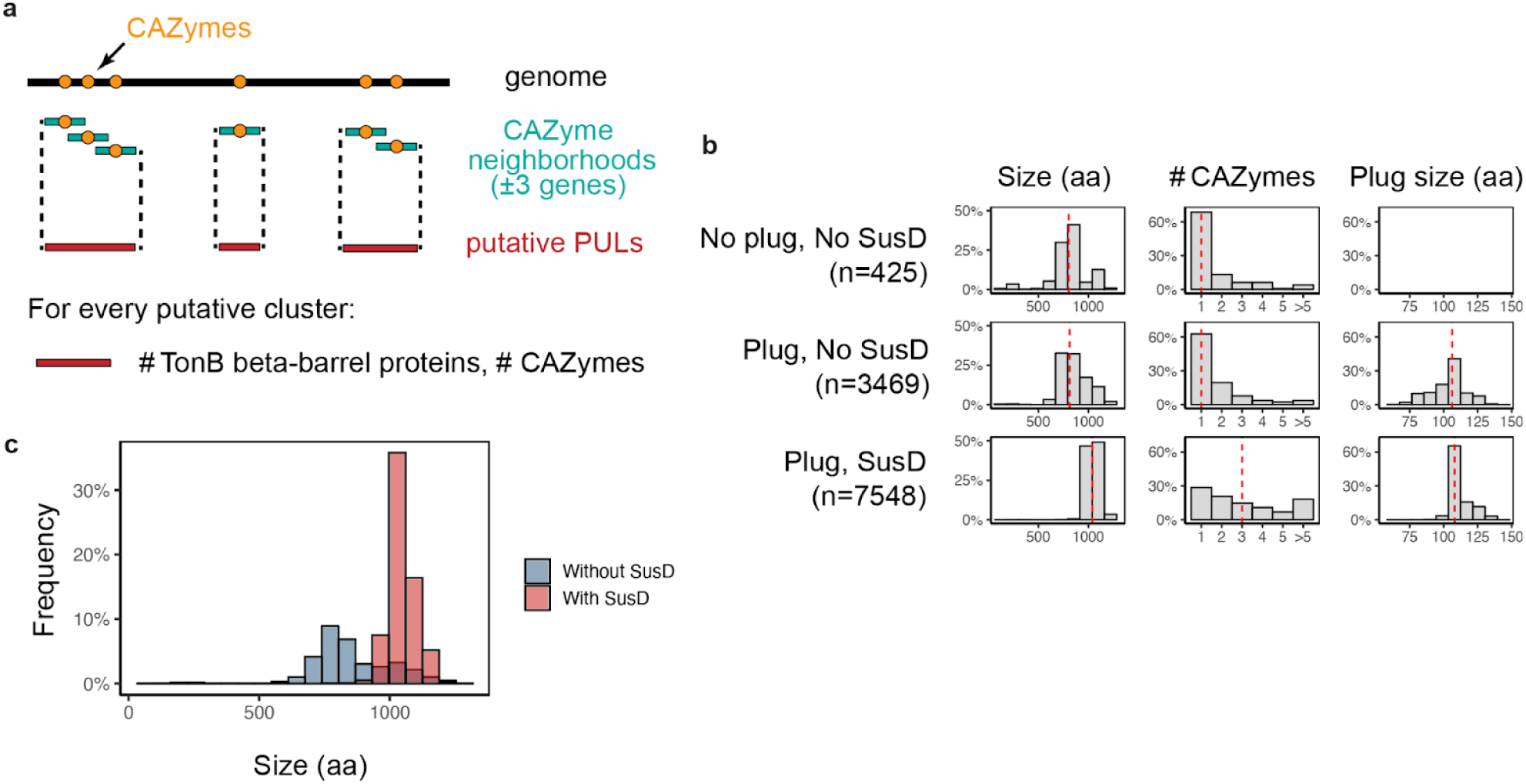
a) Schematic of the algorithm used to identify putative glycan utilization clusters not associated with SusCD pairs. All available reference genomes of scaffold quality or higher from the Bacteroidota phylum were downloaded from NCBI (n = 1496). For each genome, CAZymes from the GH, PL, CE, AA, and CBM families were annotated and their genomic neighborhoods were identified (±3 genes). Overlapping neighborhoods were merged to capture the modular organization of polysaccharide-degrading loci. Candidate clusters were then filtered to retain those encoding a single TonB-dependent transporter, identified by the presence of a β-barrel domain (Pfam PF00593, PANTHER PTHR33367) and predicted to be secreted by SignalP. b) Distribution of TonB-dependent transporter size (in amino acids), number of neighboring CAZymes, and plug-domain length across candidate clusters. Putative clusters were classified into three structural categories based on the presence or absence of a plug domain (Pfam PF07715) and a SusD-like protein within ±2 genes of the TonB receptor. The number of proteins identified in each category is shown in parentheses, and red dashed lines indicate the median value for each distribution. c) Distribution of TonB-dependent transporter size as a function of the presence of a SusD lid. Histogram shows the relative frequency of transporter lengths for clusters containing a SusD-like protein (red) or lacking one (blue). Transporters associated with SusD are generally larger, consistent with their specialization in the uptake of complex glycans.

**Figure S7.**
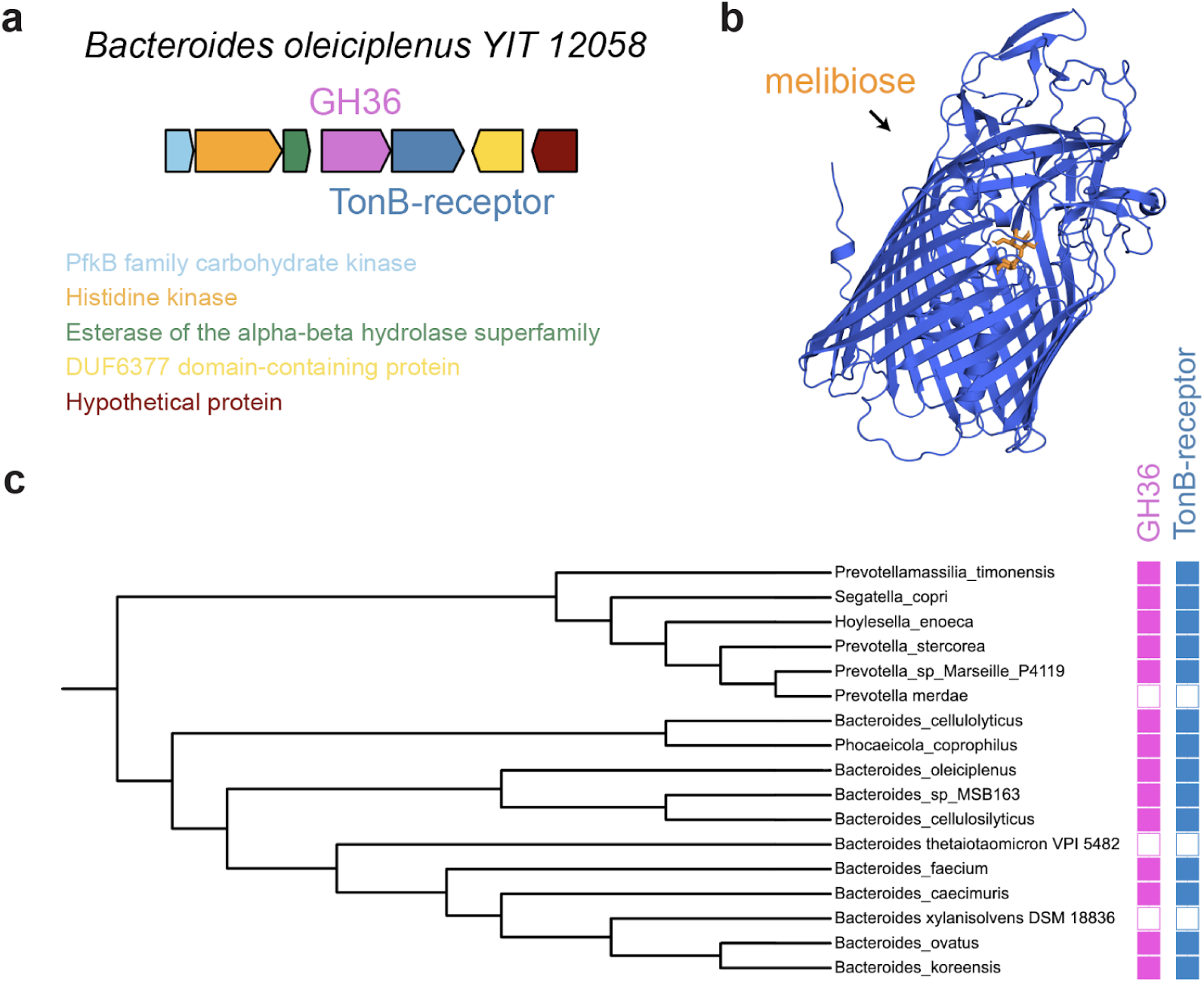
A putative Type S melibiose/raffinose transporter in the Bacteroidetes. a) Glycan utilization gene cluster from *Bacteroides oleiciplenus* identified through the bioinformatic analysis described in Fig. S6. The locus contains a Type S TonB-dependent transporter lacking both a SusD-like lid and a plug domain, adjacent to a GH36 enzyme annotated by dbCAN as EC 3.2.1.22 (α-galactosidase). The cluster also includes genes encoding a histidine and a carbohydrate kinase, forming a putative carbohydrate-utilization locus. This genomic configuration suggests a role in the uptake and hydrolysis of small α-galactosides such as melibiose or raffinose, motivating the docking simulations shown in b. b) Docking of melibiose to the putative Type S transporter revealed a binding pocket with Vina scores of –7.0 to –7.3 kcal mol⁻¹. Docking of raffinose converged on the same cavity (–6.9 to –7.9 kcal mol⁻¹) and engaged overlapping residues, indicating that the transporter can in principle accommodate both disaccharide and trisaccharide substrates. c) Homologous transporters identified across the Bacteroidetes phylum (DIAMOND ≥ 60 % amino-acid identity, ≥ 70 % coverage) were scored for adjacency to GH36 genes within ± 2 genes. Co-localization with GH36 α-galactosidases was observed in all the homologous transporters, across species from different genera, supporting a role for these genes in the uptake of α-galactoside molecules. A few type strains were included in the phylogenetic tree for reference.

